# Thermal conditioning of quail embryos has transgenerational and reversible long-term effects

**DOI:** 10.1101/2023.02.02.526800

**Authors:** Anaïs Vitorino Carvalho, Christelle Hennequet-Antier, Romuald Rouger, Joël Delaveau, Thierry Bordeau, Sabine Crochet, Nathalie Couroussé, Frederique Pitel, Anne Collin, Vincent Coustham

## Abstract

In the current context of global warming, thermal manipulation of avian embryos has received increasing attention as a strategy to promote heat tolerance in avian species by simply increasing the egg incubation temperature. However, because of their likely epigenetic origin, thermal manipulation effects may last more than one generation with consequences for the poultry industry. We found that thermal manipulation repeated during 4 generations had an effect on hatchability, body weight, and weight of eggs laid in Japanese quails, with some effects increasing in importance over generations. Moreover, we show that the effects on body weight and egg weight can be transmitted transgenerationally, suggesting non-genetic inheritance mechanisms. This hypothesis is reinforced by the observed reversion of the effect on growth after five unexposed generations. Interestingly, we demonstrate a non-transgenerational effect of thermal manipulation on heat tolerance few days after hatch. In conclusion, our multigenerational study suggests a quantitative, transgenerational and reversible effect of thermal manipulation.

## Introduction

The embryonic development is a period of environmental sensitivity that allows for phenotypic programming^1^. Notably, thermal manipulation (TM) during bird embryogenesis, a method that involves exposing the avian embryo to cyclic increases in temperature during incubation of the egg, was successfully used to improve thermal tolerance of birds during the post-hatch life^2^. In broiler chickens, an increase in incubation temperature of 1.7°C, 12 hours per day, from day 7 to 16 of incubation, was shown to reduce mortality by half in males when exposed to a heat challenge at 35 days of life^3^. Since then, a large number of TM studies have been reported in the literature in different avian species^4^, and such strategies have also been employed in fish^5^. TM was recently transposed to Japanese quails with a similar increase in incubation temperature from days 0 to 13 of incubation^6^. TM in quails had an impact on survival and growth as well as physiological and metabolic changes that suggested thermoregulatory changes. Thermal tolerance was assessed by a heat challenge at 36°C for 7 hours at 35 days of age, and while TM quails responded differently to the heat challenge compared to controls on some criteria, no clear differences in heat tolerance were observed. Similarly to what was observed in chickens^7^, these changes were associated with changes in gene expression in the TM group exposed to heat as shown in a transcriptome study^8^, suggesting gene response programming by TM.

So far, the molecular mechanisms behind gene expression changes have only been investigated in chickens. Although the mechanisms are not yet fully understood, an epigenetic mechanism involving both H3K4me3 and H3K27me3 histone marks in hypothalamus was shown to be associated to differential gene expression due to the embryonic conditioning of temperature tolerance^9^. Given the function of neighboring genes, the study showed that TM-induced epigenetic reprogramming is likely to affect the expression of genes controlling thermoregulation-related metabolic processes. In some cases, it has been shown that environmentally-induced epigenetic changes can be transmitted from one generation to the next^10,11^. Moreover, acute heat exposure of the shrimp *Artemia* influenced multiple thermal tolerance traits across generations^12^. Therefore, TM effects may last longer than the treated generation, especially if treatment is applied for several generations consecutively as a quantitative relationship may exist between the duration of the stimulus exposure and the epigenetic response^13^. Interestingly, a study comparing epigenetic lines induced by the injection of the endocrine disruptor genistein showed that the embryonic environment of quail affects the phenotype of offspring three generations later, possibly through epigenetic effects^14^. Here, we took advantage of the embryonic TM model to explore the multigenerational effects and transgenerational transmission of environmentally induced phenotypes in Japanese quails. By focusing on multiple phenotype criteria, we assessed the quantitative nature of the embryonic thermal exposure, the transgenerational inheritance of traits and the reversibility of the effects as, per nature, epigenetics effects are reversible as non-fixed within the genetic code.

## Results

### TM had multigenerational effects on body weight and on the weight of laid eggs

TM was applied on quail eggs for four consecutive generations in parallel to four consecutive generations of control incubation performed in a mirror manner (Figure 1). A comprehensive phenotypic analysis was performed at all generations and physiological parameters were analyzed at the 4^th^ generation. TM had a negative effect on the number of eggs hatched, associated with an increase in the rate of dead embryos in the shell from G2 to G4 or that failed to peck their way out of the shell from G2 to G3 (Supplementary Table 2). TM had no effect on post-hatch mortality of animals during the first 4 weeks of life, sex ratio nor body temperature at 35 days of life in all generations (Supplementary Tables 2-4). As some blood parameters were affected by TM in chickens^15^ but not quails^6^ after only one generation of TM, we performed an analysis of blood parameters at D35 on the 4th generation of animals to see if significant changes could be detected thanks to the repeated embryonic treatment over generations. However, no effect of TM was observed on blood gases, electrolytes, and metabolites compared to animals incubated in control conditions (TM4 *vs* C4; Supplementary Table 5, “multig. analysis” columns).

**Figure 1:**
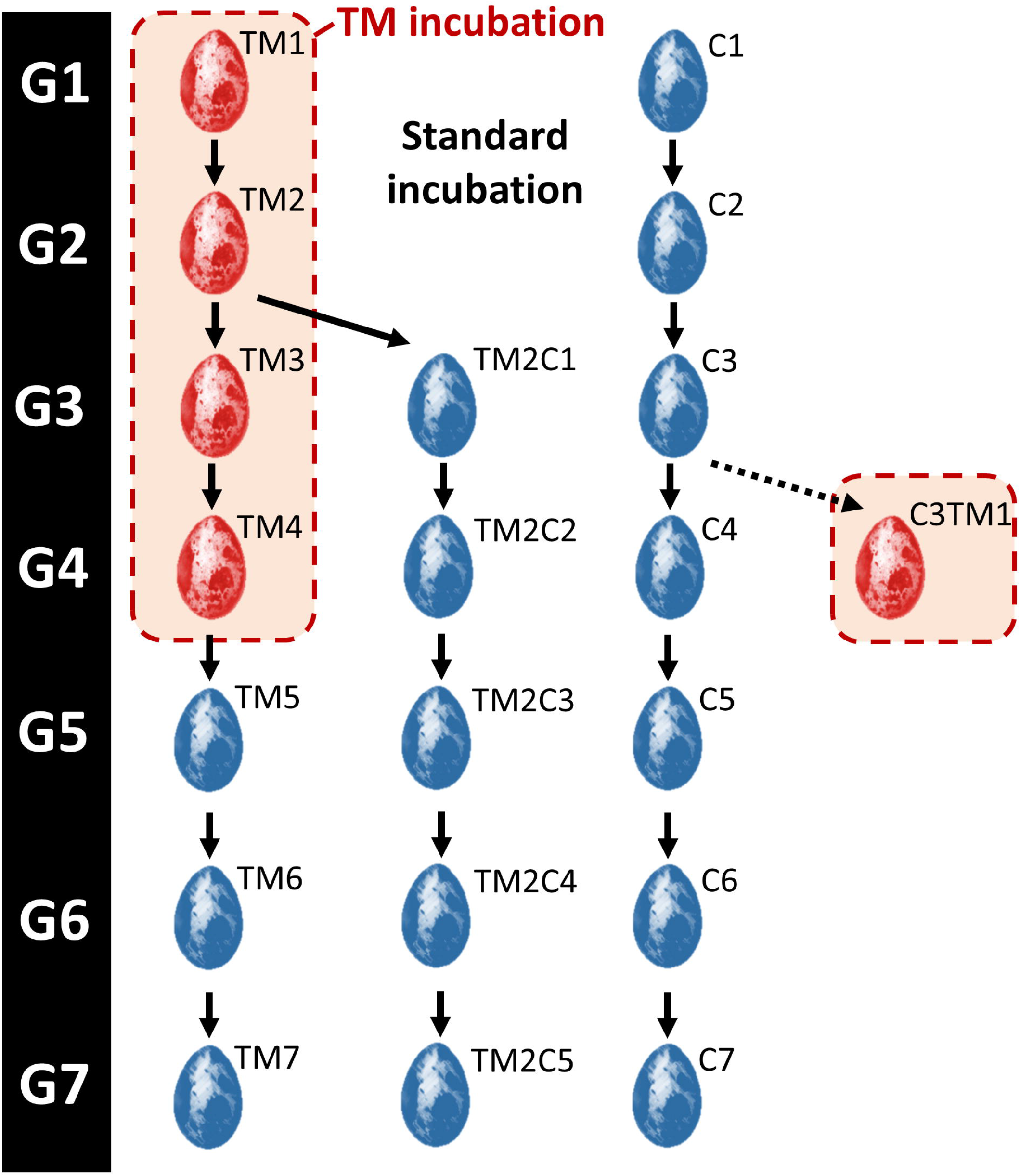
Schematic representation of the multigenerational experience plan. Eggs incubated in standard control conditions (37.8°C) are shown in blue. Thermo-manipulated eggs (TM; +1.7 °C 12h / day from incubation days 0-13) are shown in red. The name of the treatments for each generation is indicated on the top right side of the egg as follows: C*n*: *n* generations incubated in control conditions; TM*n*: *n* consecutive generations of TM; TM2C*n-2*: 2 consecutive generations of TM followed by *n*-2 generations incubated in control conditions. G*n*: Generation *n*.

TM had varying effects on quail body weight at hatch (D1), with a positive effect in the 1^st^ generation, no effect on body weight in the 2^nd^ and 3^rd^ generations, and a negative effect on body weight in the 4^th^ generation (Figure 2B and Supplementary Table 2). However, at 4 weeks of life, TM had a consistent negative effect on body weight over the 4 generations except for the females of the first generation (Figure 2A and C, Supplementary Table 2). Interestingly, at 5 weeks of life, TM had no significant effect on quail weight in the first two generations, as previously reported^6^, but the effect was significant from the 3^rd^ consecutive generation of TM (Figure 2A and D, Supplementary Table 2). This result suggests that the effects of TM on growth last longer during development as the number of generations of exposure increases.

**Figure 2:**
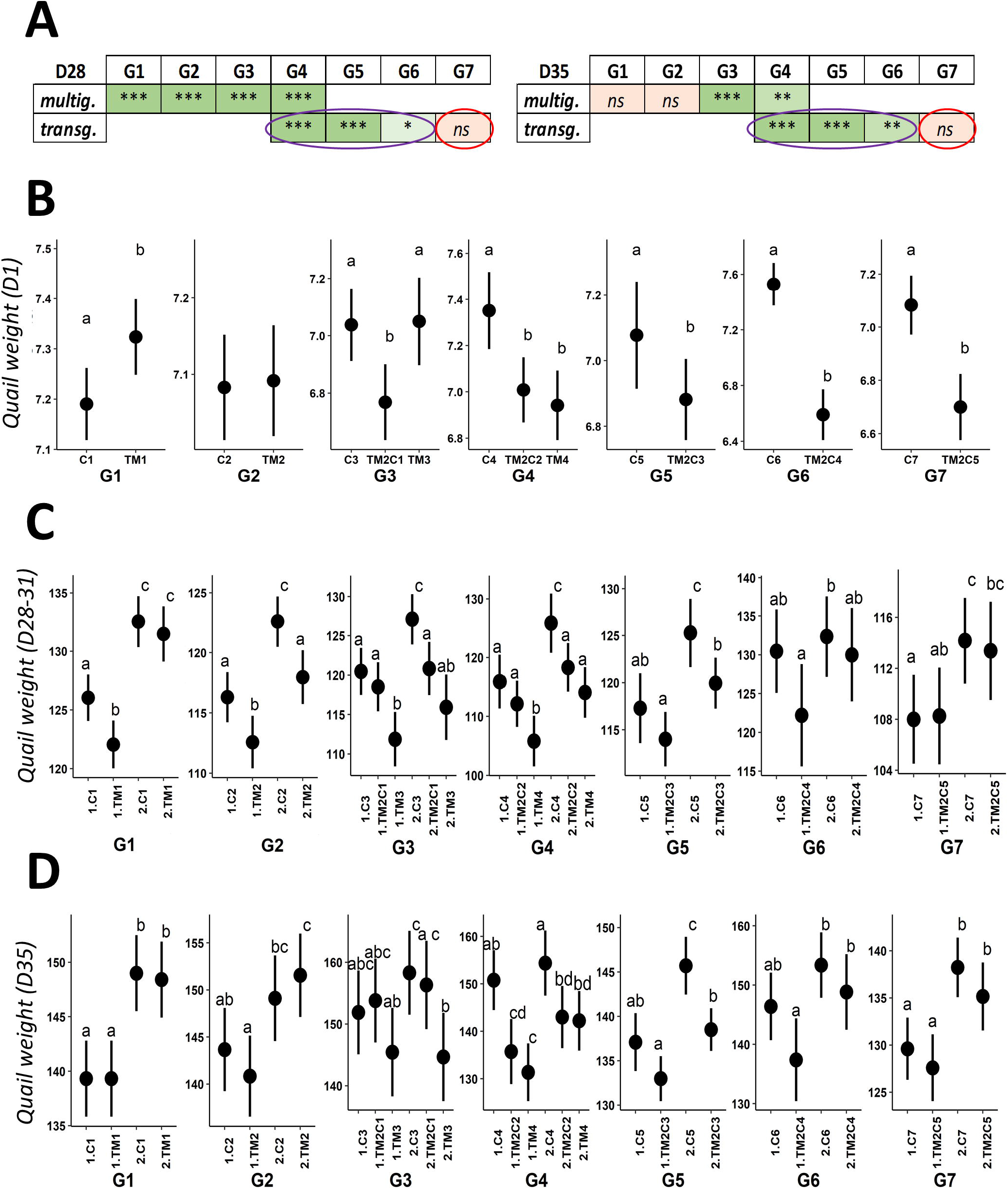
TM affects the growth of hatched quails. **A**. Summary of the statistical results of weight analyses at 4 weeks of age (left) and 5 weeks of age (right). For G1, the 4 weeks measurement has been performed at D31 instead of D28 for all other generations. Complete results including p-values and the effects on the sex and the interaction between treatment and sex can be found in Supplementary Tables 2-4 and 6. For clarity, results from multigenerational and transgenerational analyses (“multig.” and “transg.”, see methods and Supplementary Table 1) are shown on two different lines. * p < 0.05; *** p < 0.001; *ns*: not significant. Color codes (light green, green and light red) reflect the significance of the results (*, *** and *ns*, respectively). The transgenerational effect of TM on weight is circled in purple and the reversion of the effect is circled in red. **B-D**. Effects of thermal manipulation on quail weight at 1 day (B), 4 weeks (C) and 5 weeks (D) of age. Mean and confidence interval were plotted by group at each generation. The weight of quails at D1 is presented independently of sex, this factor having no significant impact (Supplementary Table 2). For G1, the 4 weeks measurement has been performed at D31 instead of D28 for all other generations. C*n*: *n* generations incubated in standard control conditions; TM*n*: *n* consecutive generations of TM; TM2C*n-2*: 2 consecutive generations of TM followed by *n*-2 generations incubated in control conditions; 1: males; 2: females. Different letters indicate significant differences between groups at p < 0.05.

Concerning reproductive traits, TM had no effect on the number of eggs laid and fertility (except for G2) but had a significant effect on the weight of laid eggs from the first generation’s eggs (Figure 3A and Supplementary Table 2). Interestingly, we observed that the mean difference in weight of eggs laid by TM and C animals increased progressively from G1 to G4, with an almost tenfold difference between G1 and G4 (Figure 3B). This result also supports an increasing effect of TM as the treatment was repeated over multiple generations.

**Figure 3:**
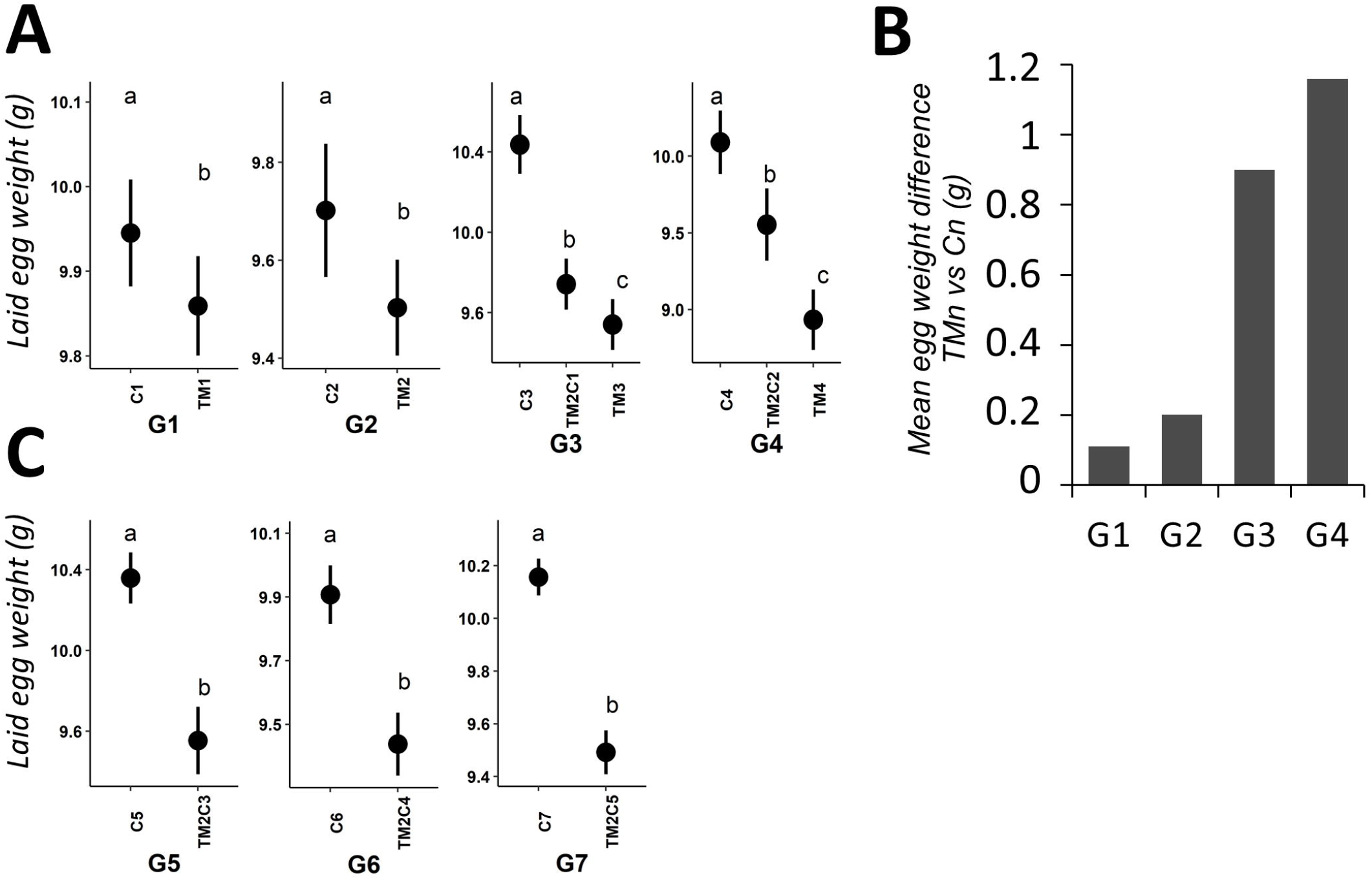
TM affects the weight of laid eggs. **A**. Effects of TM on the weight of eggs laid by G1 to G4 quails. Mean and confidence interval were plotted by group at G1-G4 generations. C*n*: *n* generations incubated in control conditions; TM*n*: *n* consecutive generations of TM; TM2C*n-2*: 2 consecutive generations of TM followed by *n*-2 generations incubated in control conditions. Different letters indicate significant differences between groups at p < 0.05. **B**. Histogram representing the mean egg weight difference laid by TM1, TM2, TM3 or TM4 quails compared to C1, C2, C3 or C4 quails, respectively. **C**. Effects of TM on the weight of eggs laid by G5 to G7 quails. C*n*: *n* generations incubated in control conditions; TM2C*n-2*: 2 consecutive generations of TM followed by *n*-2 generations incubated in control conditions. Different letters indicate significant differences between groups at p < 0.05.

### TM had transgenerational effects on body weight and on the weight of eggs laid

We next investigated whether the effects of TM on traits that were consistently affected by the treatment could be transmitted across generations or not. Given that primordial germ cells (PGCs) within the embryo are also exposed to TM, transgenerational effects of TM can only be detected from the 4^th^ generation when treatment was last applied on 2^nd^ generation embryos (effects in TM2C1 in G3 being inter-generational)^16^. Pipping defects and mortality at hatch were not transmitted to the offspring, as these effects were no longer observed when the eggs of TM parents were incubated under standard conditions (Supplementary Tables 3 and 4). However, TM had an intergenerational and a transgenerational effect on post-hatch quail weight, at D1 and both 4 and 5 weeks of age (Figure 2 and Supplementary Table 4). Sex also had an effect on weight at 4 and 5 weeks of age (females being heavier than males) but there was no interaction between sex and TM observed. A post-hoc analysis suggested that TM2C2 and TM4 animals were not different in weight compared to C4 animals in all but D28 male conditions. Remarkably, TM also had a significant intergenerational and transgenerational effect on the weight of laid eggs (Figure 3A and Supplementary Table 4). This time around, post-hoc analysis suggested that the weight of laid eggs of TM2C2 animals was intermediate to that of TM4 and C4.

### TM transgenerational effects were partly reversible

We then assessed the stability of transgenerational effects on growth and laid egg weight by maintaining the three lines (TM2C2, C4 and TM4) under standard incubation conditions for three more generations (G5 to G7). The effect of the two initial generations of TM on post-hatch weight remained significant in G5 and G6 (2^nd^ and 3^rd^ generations of transgenerational effects, respectively) but only at D1 in G7 (Figure 2). The effect on weight was indeed no longer significant in G7 at either 4 or 5 weeks of age (Figure 2A, C-D), suggesting a reversion of the effect at distance from hatching. In contrast, similarly to D1 weight observations, the effect of the two initial generations of TM on the weight of laid eggs was observed until G7 (Figure 3C), suggesting that transgenerational phenotypic stability varies according to the developmental stage and the trait observed.

### TM improved post-hatch thermal tolerance during heat exposure in a non-transmissible manner

During the present study, a heat wave has hit France in late June 2019, during the first days of rearing of the 4^th^ generation of quails. As a consequence, C4, TM4, TM2C2 and a batch of 1^st^ generation of TM animals derived from C3 parents (C3TM1) were exposed to an unexpected temperature increase of +1 to + 2.5 ° C for several hours each day, mostly between days 4 to 6 of age (Figure 4A). Unfortunately, this phenomenon induced a high mortality in the quails of the control incubation groups (C4 and TM2C1) with respectively 25.0 % (*n* = 100) and 27.4 % (*n* = 124) of dead animals during the days 3-6 period (Figure 4B). However, mortality was significantly lower in the TM incubation groups (C3TM1 and TM4) with respectively 6.7 % (*n* = 30) and 13.6 % (*n* = 103) of dead animals during the days 3-6 period (Figure 4B). This result suggests that TM improved the thermal tolerance of quails during the first days of life, but that this effect is unlikely to be transgenerational as no improvement was detected in TM2C2 animals.

**Figure 4:**
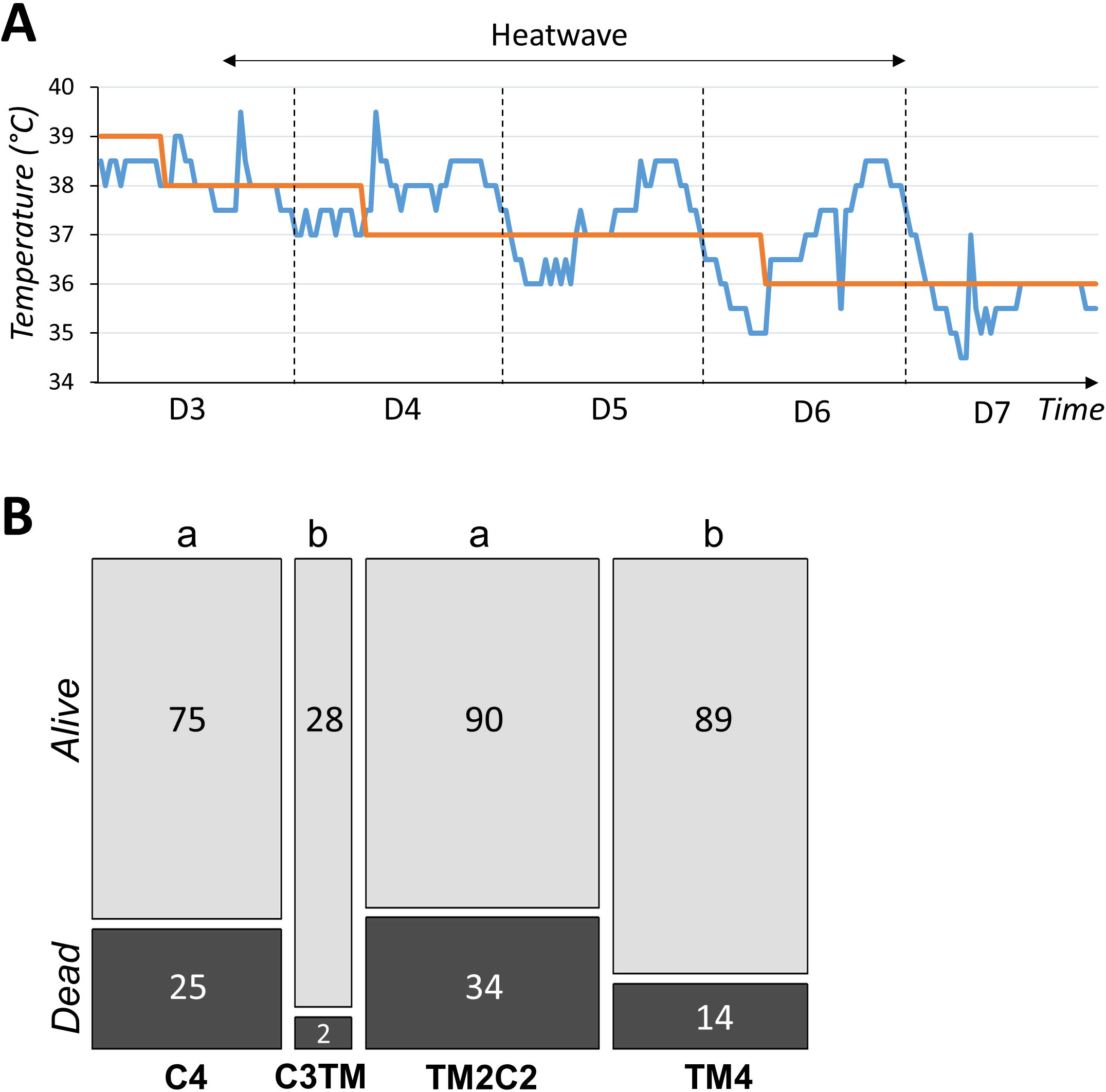
TM had a non-transgenerational and beneficial effect on heat stroke survival during the first days of life. **A**. Graph showing the theoretical set point temperature (in red) and the actual temperature (in blue) in the rearing cell as a function of time. The period of heat stress due to an uncontrolled heat wave during which mortality was measured is indicated by the double arrow. D: Day of rearing (the first day of post-hatch life being D1). **B**. Stacked histograms showing the number of animals that died or survived between days 3 and 6 of rearing during heat stroke. The width of the columns is proportional to the number of animals measured. The C3TM treatment, shown in Figure 1, corresponds to third generation control animals whose eggs were TM for the first time. C*n*: *n* generations incubated in control conditions; TM2C2: two consecutive generations of TM followed by 2 generations incubated in control conditions. Different letters indicate significant differences between groups at p < 0.05.

## Discussion

In this study, we investigated the long-term effects of TM over multiple generations. To that end, a highly inbred line of Japanese quails was used, and mirror crosses ensured that the lines would mostly vary epigenetically because of the embryonic treatment but not at the genetic level. A wide range of phenotypic measurements was collected at all generations. To our knowledge, this experiment is one of the most complete and controlled multigenerational experiment carried out on farm animals, and is of great interest for poultry farming under challenging climate conditions, which are likely to occur more frequently in the future due to the global warming context.

In the 4 generations in which TM was repeated, this study confirmed that TM has a significant impact on quail hatchability and growth as previously reported in a study that aimed at characterizing TM in quails^6^. TM had a positive effect on weight post-hatch, a negative effect at 4 weeks of age (mostly on males as revealed by the post-hoc analysis) and no effects at 5 weeks of age, in accordance with our previous results^6^. Therefore, our study confirms that the effect of TM on body weight varies during post-hatch development. Interestingly, at hatching (D1), TM no longer had a significant effect on weight in G2 and G3, and the effect on weight was significant again in G4 (and remained significant in subsequent generations despite TM was no longer applied), although this time the effect was negative, TM4 chicks being lighter than C4 at hatch. This observation must be considered in relation to the decrease of the weight of the eggs. Indeed, when comparing the percentage point differences between G1 and G4 weights, egg weights decreased by ∼10.8% while chick weights decreased by ∼8.2% over the same period. It is known that there is a significant correlation between egg weight and chick weight as well as parent age and chick weight^17^. Given that parental age was the same for all treatments and generations, at around 20-22 weeks of age, we hypothesize that egg weight may have negatively influenced chick weight over the generations. This may explain why the positive effect on D1 body weight seen in 1^st^ generation disappeared as the negative effect on egg weight seemed to increase over generations. In the two first generations, as the post-hatch development progressed, control quails were significantly heavier than TM quails at 4 weeks of age but not at 5 weeks of age. Interestingly, from the 3^rd^ generation, the negative effect of TM on weight was found both at 4 and 5 weeks of age. If a compensatory growth mechanism is involved to compensate the growth deficit of TM quails in the first two exposed generations as previously hypothesized^6^, this mechanism may no longer be sufficient as TM is repeated over generations. We believe that this phenomenon may be linked to a quantitative accumulation of epigenetic changes induced by the repetition of the treatements, as previously shown for arabidopsis plants responding to cold^13,18^. Thus, the more embryonic cells are exposed to thermal stimulus over generations, the more likely their epigenome will change state, potentially leading to longer lasting effects on the regulation of gene expression programs. This is also a first indication that some of the effects of TM can be transmitted to the offspring, otherwise no increase in effects could be observed over generations. The quantitative hypothesis is reinforced by the effect of TM on the weight of laid eggs over generations, as the observed difference seemed to increase progressively (from ∼0.1 grs in G1 to ∼1.2 grs in G4). The cumulative effect of TM was not observed for the other traits analyzed, including mortality pre- and post-hatch, sex ratio, body temperature and other reproductive parameters (number of laid eggs and fertility). Moreover, 4 generations of TM had no significant effects on blood parameters, similarly to what we previously reported for the 1^st^ generation of TM^6^, confirming previous results and suggesting that no new effect on blood parameters arose because of the repetitions of TM for 4 generations under thermoneutral conditions. Overall, TM had a significant effect on hatching, body weight, and the weight of laid eggs, and the effect on chick and egg weights seems to become more important over generations. Despite the significant differences in weight between males and females, the latter being generally the heavier, it is worth noting that both sexed generally responded in the same way to the TM treatment under thermoneutral conditions.

The main goal of the study was to investigate the potential transgenerational effects of TM in birds. To that end, given the moderate effects of TM previously reported^6^, and the quantitative epigenetic hypothesis that TM effects may be increasing over generations, we derived an epiline incubated in control conditions derived from the 2^nd^ generation of TM quails instead of the 1^st^ generation. While the deleterious effects of TM on hatching were lost upon discontinuation of treatment in G3, TM had intergenerational effects on the weight of G3 chicks at D1 and D28 but not D35, with a decrease in chick weight at hatch in TM2C1 animals compared to C3 (despite TM3 did not differ from C3, surprisingly), and an effect mainly carried by females at D28 according to the post-hoc analysis. Moreover, TM had intergenerational effects on the weight of laid eggs, the weight of TM2C1 eggs being intermediate to that of C3 and TM3. This suggests that the effects of TM related to growth were partially transmitted or memorized in the offspring. The transgenerational effect was evaluated in G4 and our analysis showed that TM had significant transgenerational effects on the post-hatch weight of quails that affected both sexes at 4 and 5 weeks of age. Similarly to what has been observed in G3, no effects on hatchability were detected in G4 either. The interpretation of a transgenerational effect on body weight was reinforced by the observation of a significant effect in G5 and G6 despite the intra-generation post-hoc analyses suggested that the differences between C5 or C6 and TM2C3 or TM2C4, respectively, were diminishing. This decrease was confirmed by a reversal of the effect on body weight observed at 4 and 5 weeks (but not at hatching) in G7. Near hatching, the transgenerational effect seemed more robust with a significant difference in D1 body weight still observable between C7 and TM2C5 animals. This observation is another argument in favor of a stronger impact of TM at a temporal proximity of the embryonic development, in agreement with the quantitative hypothesis. Moreover, the effect on the weight of laid eggs was also very robust as the reduction in egg weight induced by 2 generations of TM was still observed after 5 untreated generations. It is worth noting that, given the strong correlation between egg weight and birth weight^17^, we may not see a reversal of the effect on birth weight until the effect on egg weight is also reversed. Therefore, more generations under standard incubation conditions may be required to observe a reversal of the effects of TM on laid eggs and hatching weights.

A previous study showed that exposure during embryogenesis to the endocrine disruptor genistein had transgenerational effects on several traits including body weight in quails at 3 and 27 weeks of age, probably through non-genetic inheritance mechanisms^14^. Here, the combination of several lines of evidence that are the quantitative, transgenerational and reversible nature of the effects of TM is in clear agreement with the hypothesis that embryonic environmental exposures have long-term transgenerational effects under the control of non-genetic epigenetic mechanisms. Indeed, reversion would probably not have been observed if the effect was caused by genetic variants. Furthermore, the risk of genetic differences between lines was unlikely given our mirror-crossing strategy and the high level of inbreeding of the quails used. The transient nature of the effects is a characteristic of epigenetic changes that are heritable (mitotically, and in that case also meitoically) and reversible by nature^16^. Epigenetic alterations would then translate into gene expression changes leading to phenotypic alterations. We previously showed that TM has an impact on the transcriptome of quails in response to a 7-hours 36 °C heat challenge at 5 weeks of age^8^. Interestingly, the effect on gene expression was mainly on females, which could explain why the transgenerational effect on body weight is mostly affecting females at D28 and D35 in G4 and G5 according to the post-hoc analysis. Investigation of the epigenome of C4, TM4 and TM2C2 would be required to confirm the epigenetic nature of TM conditioning on gene expression.

TM is an embryonic phenotype programming strategy that was initially developed to improve post-hatch heat tolerance, as shown for 35-day-old male broilers that better tolerated an acute heat challenge^3^. However, we did not observe such improvement in quails in a heat exposure at the same age^6^. At the start of the 4^th^ generation, an unexpected heat wave led to an increase in rearing temperature conditions beyond the recommended values, with increases of 1.0 to 2.5 °C above the set values for several hours despite our efforts to keep this increase as limited as possible. These temperature rises occurred when the quail’s thermoregulatory system was probably not mature yet, as it sets up around days 3 to 5 in chickens^19^. This may explain why excess mortality was observed during the heat stroke. However, the distribution of mortalities between groups showed an effect of TM, with a significant increase in the mortality of animals that have not been directly treated by TM during the 4^th^ generation of incubation (C4, TM2C2). In addition, we incorporated a “first generation” TM treatment (TM1) derived from C3 parents. Interestingly, these animals responded in the same way as the 4th generation TM animals (TM4) with a lower mortality rate than the standard incubated animals. These results suggest that TM may improve the thermal tolerance of quail during the first days of life, regardless of the generation exposed, which could be related to the immaturity of the quail’s thermoregulatory system at start-up. This would be the first demonstration of a beneficial effect of thermal manipulation on heat tolerance in this species. However, this positive effect is not likely to be transmitted to the offspring, as no improvement of survival was observed in TM2C2 animals compared to C4 quails.

In conclusion, TM has a wide range of effects that can be transmitted (body weight, egg weight) or not (mortality, thermal tolerance) to the offspring. These effects appeared to vary in intensity according to the trait observed, the developmental stage and the sex. While the effect on temperature tolerance was beneficial as TM improved survival when animals were heat-challenged during the first days of post-hatch life, many of the effects of the embryonic thermal protocol were detrimental as it decreased hatchability and lowered growth performance, in particular when the treatment was repeated across generations. As previously proposed^6^, TM procedure should be refined to mitigate deleterious effects while keeping tolerance advantages. For example, this may be achieved by reducing the duration of TM that encompasses very early stages of development, as the treatment starts from the 12^th^ hour of incubation. In any case, should TM become a common practice in hatcheries, this treatment could be advantageously used on terminal products reared in hot climates or during summers (the effects on growth fading away after hatching) but it should be used sparingly on breeders as new phenotypes may appear over generations. This is not only relevant for poultry farming, but can also be of interest for other ectothermic species, as thermal acclimation capacity can be modified by temperature in the early development of fishes^5^. In general, this work teaches us that any phenotype conditioning procedure in livestock should be carefully studied over several generations to ensure that these new practices do not cause unexpected health or performance issues later on.

## Methods

### Incubation and rearing conditions

All experiments were carried out in accordance with the legislation governing the ethical treatment of birds and were approved by the French Ministry of Higher Education and the Val-de-Loire Animal Ethics Committee (authorization N° APAFIS #4606–2016032111363124).

Animal rearing was performed in the PEAT INRAE Poultry Experimental Facility (2018, https://doi.org/10.15454/1.5572326250887292E12).

The highly inbred INRAE Cons DD Japanese quail line was used in this study as previously described^6^. Eggs were collected during 14 days from mothers of similar age at each generation (about 20-22 weeks of age) and stored at 12°C before incubation. Control eggs (C) were incubated at 37.8°C and 56% relative humidity (RH) during the whole incubation period in an automated commercial incubator with regulation of temperature, humidity and ventilation (Bekoto B64-S, Pont-Saint-Martin, France). Thermal manipulation (TM) of 39.5°C and 65% RH was applied for 12 h/d from I0 (from the 12^th^ hour of incubation) to I13, in a second identical incubator. The incubation of C eggs began 12 h before the incubation of TM eggs to synchronize hatching, 12 h corresponding to the acceleration of embryonic development induced by heat^6^. All eggs were subjected to a turning angle of 90 degrees every hour. At I14, the presence of living embryos was checked by candling and eggs were transferred to one hatcher and incubated at 37.5°C with RH between 75 and 80 % until I18. Hatched quails were weighed and identified with a numbered wing tag at I17 (peak hatch rate, also considered the beginning of out-of-shell life). Then, the quails were transferred to a single concrete-floored room covered with litter bedding and reared under standard conditions: temperature was progressively reduced from 39-40°C on the first day of rearing D1 to 20°C on D24, and water and feed were supplied *ad libitum*. Ambient temperature in the rearing room was recorded with a PLUG&TRACK DAL0075 22Lx thermo-button located near the drinking ramp at quail’s height.

### Multigenerational design

The breeding strategy was defined in order to limit genetic divergence between the three lines kept under the different treatments. For the base generation (G0), eggs produced by 73 founder couples were equally dispatched between treatments (C and TM) to obtain homogeneous G1 egg groups: 480 eggs with an average weight of 10.16 g ± 0.76 for the C incubation group and 481 eggs with an average weight of 10.17 g ± 0.72 for the TM incubation group. For subsequent generations (G2, G3 and G4), and when this was possible, divergence between lines was constrained by directing the mating in order to maximise co-ancestry between individuals across treatments (i.e. the number of common ancestors, Supplementary Figure 1). Briefly, 19 TM and C mirror couples were used to produce G2 eggs. 23 C2 and 46 TM2 mirror couples were used to produce G3 eggs (TM2 couples were doubled to produce TM3 and TM2C1 eggs). Finally, 8 C3, TM3 and TM2C1 mirror couples were used to produce G4 eggs that yield the 4^th^ generation animals, i.e., C4, TM4 and TM2C2.

This breeding strategy was relaxed for the production of generations 5 and 6 where mating within each line were decided in order to minimise average inbreeding. An optimisation program using a simulated annealing approach was used for this purpose (SYSAAF, program available on request). The pedigree of each individual was known 3 generations before the base generation. Divergence between populations was measured using pairwise F_ST_ values between populations using pedigree information as described by Caballero and Toro (2002). Efficiency of the breeding strategy to limit genetic divergence between lines was verified by comparing the observed pairwise F_ST_ values to the 95% confidence interval obtained after 1000 simulations where pedigree was randomised within each line (Supplementary Figure 2). Overall, the breeding strategy permitted to keep low levels of divergence between lines for generation 1 to generation 4. The increased divergence at generation 5 is due to a strong bottleneck within each treatment.

### Zootechnical measurements

Embryonic development parameters, such as proportion of embryos that died in-shell (eggs without evidence of pipping) or embryos with external pipping (eggs not fully hatched), were assessed at I18 Each day of rearing, mortality was recorded and early mortality was assessed at D28. At D28 or D31, all animals were visually sexed, and the sex ratio was estimated among live animals. For the remainder of the analysis, only quails with a determined phenotypic sex and a wing identification tag at D35 were included in the statistical analysis from the collected data. At D1, D28 or D31, and D35, all quails or a random selection of quails were weighed. At D35, internal temperatures were measured using a KIMO KTT-310 probe thermometer inserted in the cloaca of a randomly selected set of animals. In G4, blood samples were collected once from a randomly selected subset of quails for hematological analyses. Sample sizes are shown in Supplementary Tables 2-4 and 6. Egg-laying parameters were collected during 65 days from the pairing. Fertility rate was determined as the number of eggs with developed embryos observed by candling at I14 relative to the total number of incubated eggs.

### Hematological analyses

One mL of blood was sampled from a randomly selected subset of G4 quail using a heparinized needle and syringe (SS02SE1, Terumo, Guyancourt, France; BD Microlance™ 3 23G, BD, Le Pont-De-Claix, France). Blood gases and electrolytes were assessed using an IRMA True Point blood analysis system and single-use CC cartridges (ITC Nexus DI, Edison, NJ). Partial pressure of carbon dioxide (pCO2) and oxygen (pO2), pH, hematocrit (Hct), sodium (Na+), potassium (K+), ion calcium (iCa), bicarbonate (HCO3-), total carbon dioxide (TCO2) concentrations, base excess in blood (Beb), base excess in extra-cellular fluid (Beecf), oxygen saturation percentage (O2sat), and total hemoglobin (tHb) were measured.

### Statistical analyses

Statistical analyses were performed with R software^21^ version 4.2.1 using the packages base, lsmeans^22^ version 2.30-0 and multcomp^23^ version 1.4-20. Chi-square tests of homogeneity and hypothesis tests in the linear models were performed at a 5 % statistical significance level. All statistical parameters including tests used, variables involved in tests, specific comparisons of treatments and sample size are summarized in Supplementary Table 1. Briefly, three types of analysis were performed: multigenerational analysis (“multig.”) comparing only TM and C treatments (from G1 to G4); intergenerational analysis (“interg.”) only in G3 comparing C3, TM2C1 and TM3 treatments; and transgenerational analysis (“transg.”) comparing C4-7, TM4-7 and TM2C2-5 conditions from G4 to G7 (Supplementary Table 1). After a statistically significant result with a linear model, post-hoc analyses were performed with Tukey’s multiple comparisons adjustment.

## Data availability

The authors declare that all data supporting the findings in this study are available within the article and its supplementary information files. Raw phenotypic datasets are available from the *Recherche Data Gouv* public repository entitle “Multigenerational analysis of thermal manipulation of Japanese quail”, DOI: https://doi.org/10.57745/D38TFS.

## Acknowledgements

This study was funded by the ANR JCJC “QuailHeatE” Research Program [ANR-15-CE02-0009-01]. We are grateful to all the members of the MOQA team at INRAE for assistance with sample collection and to the staff of the INRAE PEAT (2018, https://doi.org/10.15454/1.5572326250887292E12) Poultry Experimental Facility for animal care and management. The authors thank Christine Leterrier and Ludovic Calandreau for their support during the transgenerational rearing of animals.

## Authors’ contributions

V.C., A.V.C., J.D., R.R., F.P. and A.C. designed the experiments. A.V.C., V.C., J.D., S.C., T.B. and N.C. performed the experiments. V.C., A.V.C., C.H.-A. and R.R. analyzed the results. V.C., A.V.C. and R.R. wrote the manuscript. All authors read and approved the final manuscript.

