## Supplementary Information for "Thermal conditioning of quail embryos has transgenerational and reversible long-term effects"

**Legends for Supplementary Tables**

**Supplementary Table 1: Statistical parameters for the transgenerational analysis** (variables, sample sizes and statistical models). Lm: Linear model. 1= males. 2= females. C*n*= *n* generations incubated in control conditions. TM*n*= *n* consecutive TM generations. -= non applicable statistical test. ND= not determined. The sample sizes are indicated in parentheses.

**Supplementary Table 2: Multigenerational analysis results**. C*n*= *n* consecutive control generations. TM*n*= *n* generations incubated in control conditions. Significant results are highlighted in bold.

**Supplementary Table 3: Intergenerational analysis results**. C*n*= *n* consecutive control generations. TM*n*= *n* generations incubated in control conditions. Significant results are highlighted in bold.

**Supplementary Table 4: Transgenerational analysis results**. C*n*= *n* consecutive control generations. TM*n*= *n* generations incubated in control conditions. Significant results are highlighted in bold.

**Supplementary Table 5: Blood parameters analysis results (G4)**. C*n*= *n* consecutive control generations. TM*n*= *n* generations incubated in control conditions. Significant results are highlighted in bold.

**Supplementary Table 6: G5-G7 results**. C*n*= *n* consecutive control generations. TM*n*= *n* generations incubated in control conditions. Significant results are highlighted in bold.

**Supplementary Figure 1:** **Description of the breeding strategy**. Males and females are represented by green and red circles respectively. Letters in the base generation symbolize different individuals; letters in subsequent generations are the base population ancestors of each individual. Families produced by mating in the base generation were split between the two treatments TM and C. Mating among individuals in the first generation were organized in order to maximize the number of common ancestors between individuals across the two treatments in the second generation. A similar breeding strategy was used to produce generations 3 and 4.

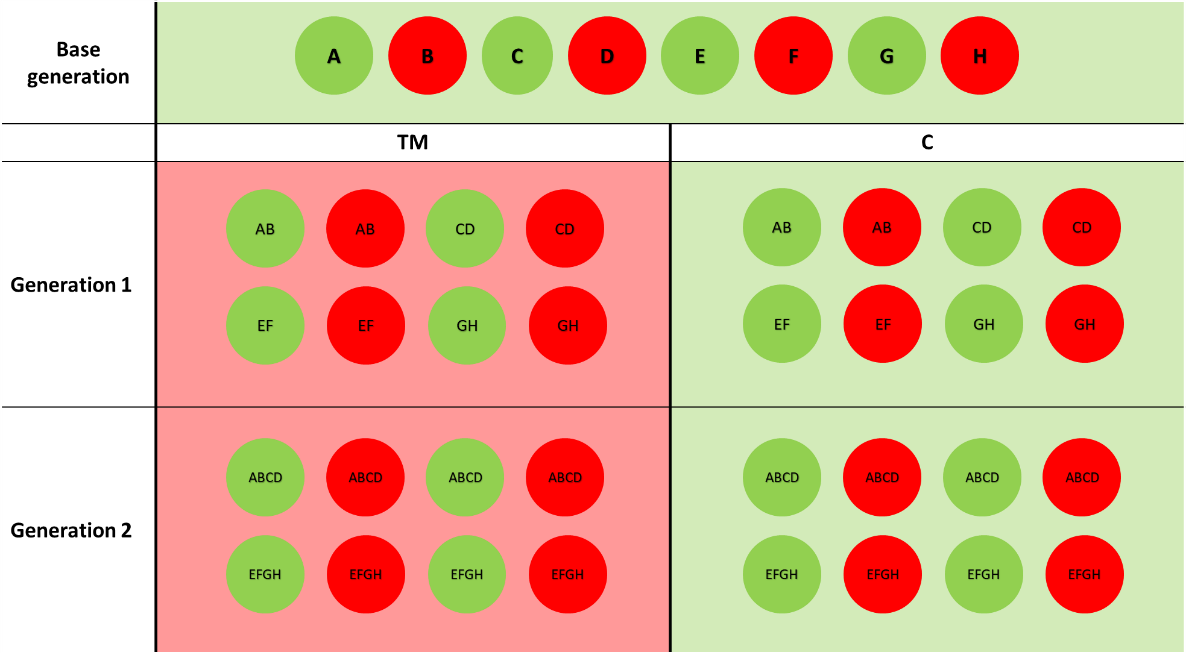

**Supplementary Figure 2: Pairwise Fst values between lines**. Red lines represent the 95% confidence interval obtained after 1000 simulations where pedigree was randomized within each line.

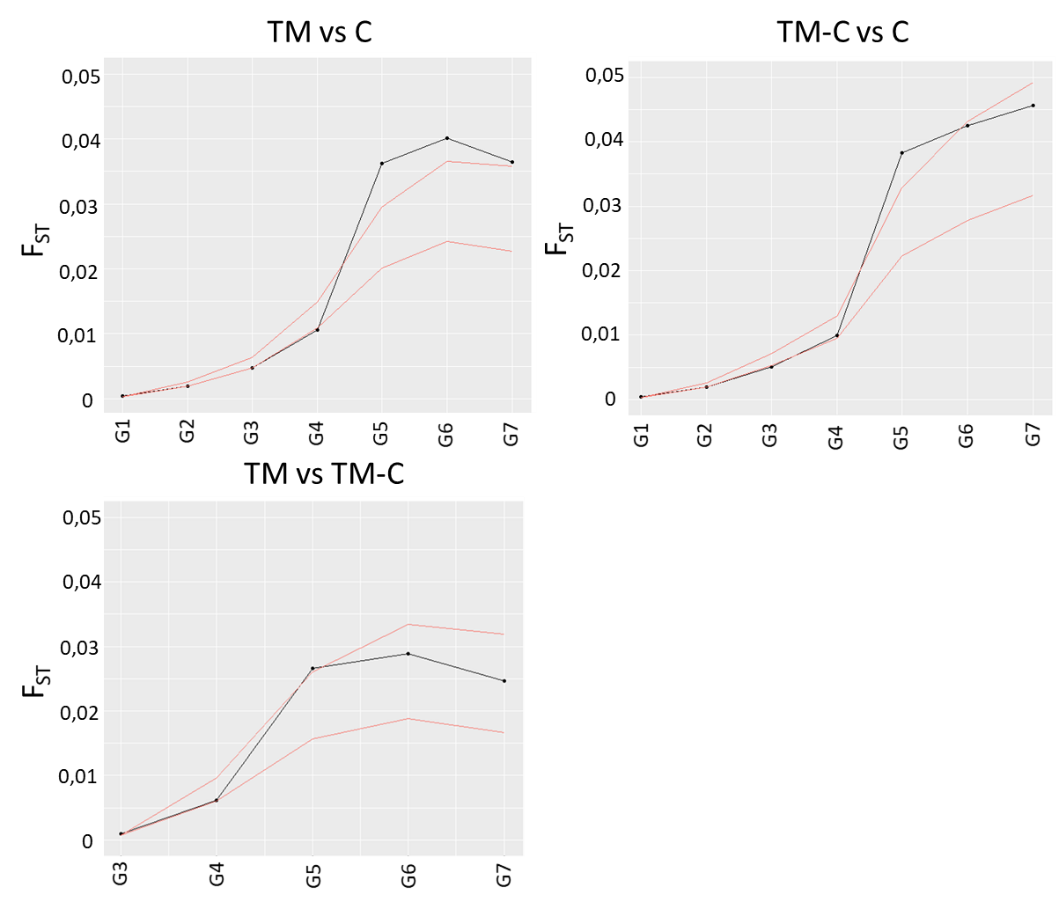
